# Mucin 4 Protects Female Mice from Coronavirus Pathogenesis

**DOI:** 10.1101/2020.02.19.957118

**Authors:** Jessica A. Plante, Kenneth S. Plante, Lisa E. Gralinski, Anne Beall, Martin T. Ferris, Daniel Bottomly, Richard Green, Shannon K. McWeeney, Mark T. Heise, Ralph S. Baric, Vineet D. Menachery

## Abstract

Using incipient lines of the Collaborative Cross (CC), a murine genetic reference population, we previously identified a quantitative trait loci (QTL) associated with low SARS-CoV titer. In this study, we integrated sequence information and RNA expression of genes within the QTL to identify mucin 4 (*Muc4*) as a high priority candidate for controlling SARS-CoV titer in the lung. To test this hypothesis, we infected *Muc4*^-/-^ mice and found that female, but not male, *Muc4*^*-/-*^ mice developed more weight loss and disease following infection with SARS-CoV. Female *Muc4*^*-/-*^ mice also had more difficulty breathing despite reduced lung pathology; however, no change in viral titers was observed. Comparing across viral families, studies with chikungunya virus, a mosquito-borne arthralgic virus, suggests that Muc4’s impact on viral pathogenesis may be widespread. Although not confirming the original titer QTL, our data identifies a role for Muc4 in the SARS-CoV disease and viral pathogenesis.

**Importance:** Given the recent emergence of SARS-CoV-2, this work suggest that *Muc4* expression plays a protective role in female mice not conserved in male mice following SARS-CoV infection. With the SARS-CoV-2 outbreak continuing, treatments that modulate or enhance *Muc4* activity may provide an avenue for treatment and improved outcomes. In addition, the work highlights the importance of studying host factors including host genetics and biological sex as key parameters influencing infection and disease outcomes.

## Introduction

Most viral infections present with an array of symptoms following human infection, ranging from asymptomatic to self-limiting, chronic, and sometimes fatal disease. Some of this diversity is mediated by age, sex, and other comorbidities, but these demographics do not account for all of the variability in disease outcomes. Genome-wide association studies (GWAS) and candidate gene studies have established a number of genes associated with human susceptibility to infectious diseases, but are limited by the large populations required to detect genetic effects and by existing knowledge gaps (Fellay et al., 2007; Ge et al., 2009; Lindesmith et al., 2003; López et al., 2010). Furthermore, during outbreak settings there is often limited access to clinical samples from infected humans. Thus, host susceptibility allele identification is oftentimes heavily compromised by situational expediency and new approaches are needed to identify alleles that regulate emerging virus pathogenesis.

Severe acute respiratory syndrome virus (SARS-CoV) is a respiratory pathogen that first emerged in the Guangdong Province of China in November 2002, and rapidly spread to 28 countries, resulting in over 8,000 cases with a 10% case fatality ratio. The subsequent emergence of Middle East respiratory syndrome coronavirus (MERS-CoV) and the discovery of several other related “pre-epidemic” bat coronaviruses that seem poised for sudden emergence emphasize the need for a comprehensive understanding of how highly pathogenic human coronaviruses, such as SARS-CoV, interact with their hosts to regulate disease severity (Coleman and Frieman, 2013; Menachery et al., 2015b; Menachery et al., 2016). The importance of these early predictions was demonstrated by the ongoing outbreak of SARS-CoV-2 (Gorbalenya et al., 2020; Gralinski and Menachery, 2020), which has caused 45,171 confirmed cases and 1,115 deaths as of February 12, 2020 (World Health Organization, 2020).

We have developed a model of CoV infection in the genetically diverse Collaborative Cross (CC) population (Churchill et al., 2004; Collaborative Cross Consortium, 2012). This model allows us to assess the impact of host genetic variation on CoV disease, such as in the context of SARS-CoV pathogenesis (Gralinski et al., 2015). The CC panel contains mouse lines derived from a funnel breeding scheme of eight founder strains, including 5 traditional lab strains and 3 wild-derived strains (Churchill et al., 2004; Collaborative Cross Consortium, 2012; UNC Systems Genetics Core Facility, 2012). This population takes advantage of naturally-occurring genetic polymorphisms to capture approximately 90% of the common genetic diversity within *Mus musculus*, and that diversity is distributed evenly throughout the genome, with elevated minor allele frequencies compared to standard natural populations (Aylor et al., 2011; Yang et al., 2007; Yang et al., 2009; Yang et al., 2011). Our initial screen within an incompletely inbred set of CC mice (the preCC screen) (Gralinski et al., 2015), along with an F2 screen utilizing a highly resistant and highly susceptible CC mouse line (Gralinski et al., 2017), identified nine quantitative trait loci (QTLs) impacting SARS-CoV disease. Among these QTLs was locus *HrS2* on chromosome 16, which was identified as modulating SARS-CoV titer levels in the lung (Gralinski et al., 2015).

Here, we narrow the *HrS2* locus to a priority candidate gene, mucin 4 (*Muc4*), via the integration of sequence data from the CC founders (Keane et al., 2011) and RNA expression results from our preCC experiment. Low expression of *Muc4* in mice containing the PWK/PhJ allele corresponded with the low SARS-CoV titer associated with the QTL located at locus *HrS2*. We subsequently examined infection of mice lacking *Muc4*, predicting reduced viral load. However, our findings suggest that while Muc4 does not regulate SARS-CoV titer, it does have broad activity attenuating the pathogenic impact of viral infections.

## Results

### Selection of *Muc4* as a SARS-CoV disease associated gene

In earlier studies, incipient CC lines were used to identify four QTLs associated with SARS-CoV phenotypes in female mice, one of which was mapped on the basis viral titer in the lung at four DPI (Gralinski et al., 2015). A subsequent study utilizing both male and female F2 mice derived from a highly susceptible and highly resistant CC mouse line identified five more SARS-CoV-associated QTLs, three of which were mapped at least partially on the basis of viral titer (Gralinski et al., 2017). Located on chromosome 16 from nucleotides 31,583,769-36,719,997, the pre-CC QTL accounted for 22% of the variation in viral titer, which ranged from over 10^8^ PFU/lobe to below the limit of detection (10^2^ PFU/lobe). The F2 QTLs, by comparison, were located on chromosomes 18, 7, and 12 and explained 12.9%, 12.3%, and 5.4%, respectively, of the observed variation in titers from high 10^2^ to low 10^7^ PFU/lobe. Thus, while SARS-CoV titer in the lung is clearly influenced by multiple host genetic factors, the preCC QTL on chromosome 16 was chosen for candidate gene selection and validation. Analysis of allele effects revealed that the founder PWK/PhJ was the main driver of low SARS-CoV titer in the lung (Fig. 1A), and PWK/PhJ also had lower titers than any other founder strain at four DPI (Gralinski et al., 2015). We therefore used private SNPs or In/Dels found in PWK/PhJ to reduce the potential targets in the QTL to seven ncRNAs and 74 genes for downstream analysis.

**Fig. 1.**
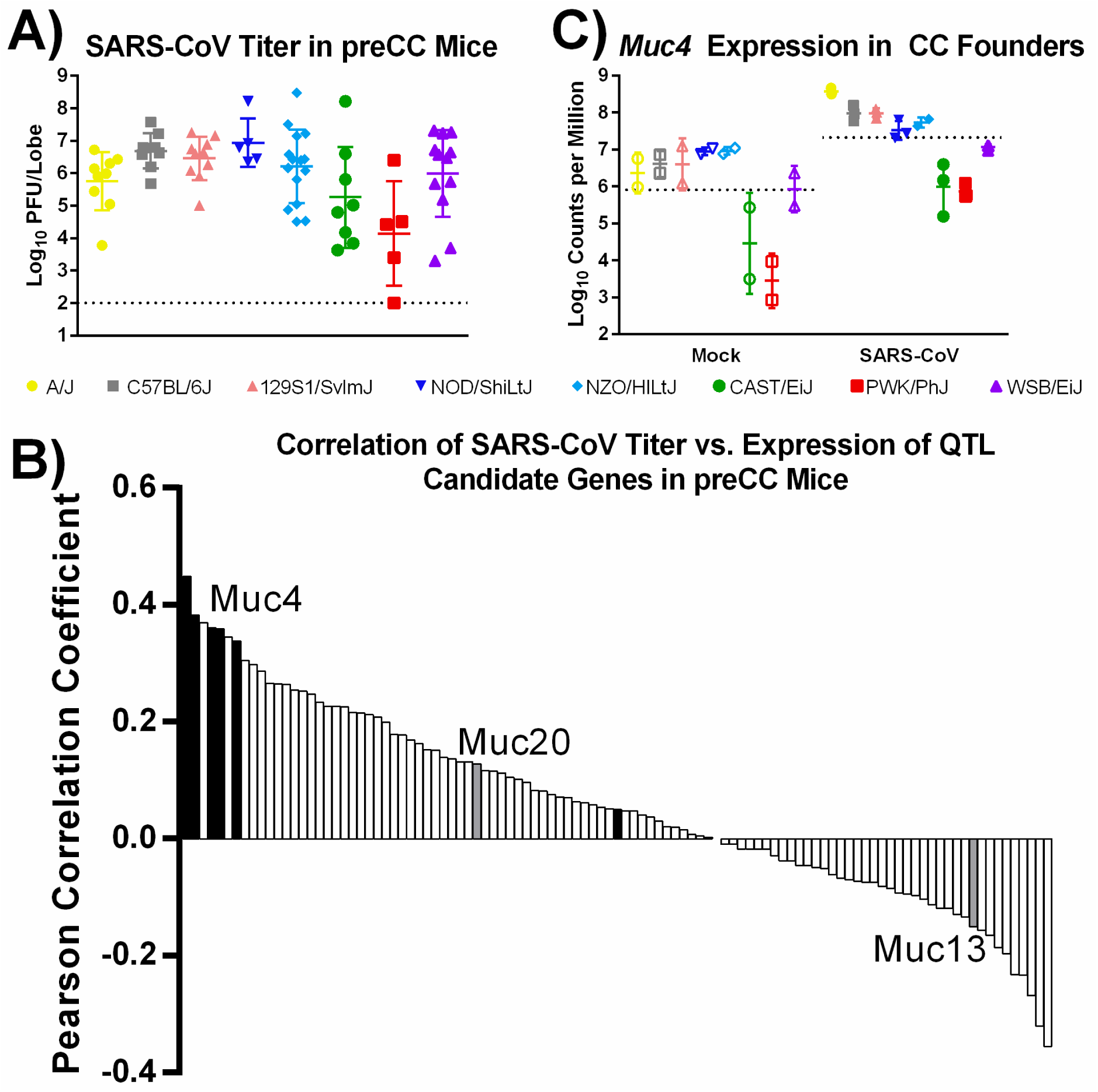
Selection of *Muc4* as a candidate gene from the preCC screen of SARS-CoV lung titer. (A) SARS-CoV lung titer at four DPI in preCC mice that were homozygous for one of the eight indicated founder strains at genotype marker nearest to the *Muc4* gene. Dashed line represents the lower limit of detection for the assay. (B) Correlation values for Log_10_-transformed viral titers and host RNA expression values as determined by microarray in the lungs of preCC mice at four DPI with SARS-CoV. Bars correspond to individual microarray probes representing genes in the titer QTL region. Black bars indicate probes for *Muc4*. Grey bars indicate probes for other mucins (*Muc13* and *Muc20*) within the QTL. Open bars represent non-mucin genes within the QTL. (C) Expression of *Muc4* RNA in the CC founder lines at four DPI following either mock (open symbols) or SARS-CoV (closed symbols). Dashed lines represent the mean expression level for all founder mice.

To prioritize the list of ncRNAs and genes within the QTL, the relationship between RNA transcript levels and viral titer in the lung was examined. Sixty seven preCC mice had both microarray and titer data available, of which sixty had titers above the limit of detection (Gralinski et al., 2015). Of the 74 candidate genes, 54 had one or more probes on the microarray (Table S4) and *Muc4* expression had the strongest correlation with viral titer, with five of the six *Muc4* probes yielding positive correlations of 0.34-0.45 (Fig. 1B, Table S5). Using infection data from the founder mice, *Muc4* RNA expression levels mimicked the allelic effects of the QTL with PWK/PhJ mice exhibiting low *Muc4* RNA levels (Fig. 1C). Other potential targets included Lrrc33, Sec22a, Parp14, and Ildr1; however, these genes either have little known linkage to viral replication and the immune response (e.g., Lrrc33, Sec22a, and Ildr1) or the directionality of the correlation did not support their relationship to viral titer in the lung (Parp14). For Muc4, high levels of expression at mucosal surfaces and in various human cancers have been reported (Andrianifahanana et al., 2001; Chaturvedi et al., 2008; Kamikawa et al., 2015). In addition, *Muc4 plays* a role in cell and anti-apoptotic signaling (Chaturvedi et al., 2008; Funes et al., 2006; Moniaux et al., 2007). Together, the data suggest that *Muc4* may play a role in viral titers at day 4 post SARS-CoV infection.

### *Muc4* drives sex-based difference in weight loss

To examine the role of *Muc4* during SARS-CoV infection, we utilized *Muc4*^*-/-*^ mice (Rowson-Hodel et al., 2018) which have no gross differences in behavior, overall health, size, or lung function, the latter phenotype measured by plethysmography. *Muc4*^*-/-*^ and WT control mice were infected with SARS-CoV and examined over a four-day time course. Following infection, a significant sex-based difference in weight loss between *Muc4*^*-/-*^ and WT mice was observed (Table S3). Female *Muc4*^*-/-*^ and control were essentially indistinguishable through two DPI (Fig. 2A). However, infected *Muc4*^-/-^ females continued to lose weight through day four, with 42% (6 of 14) meeting euthanasia requirements (≥20% weight loss) and another 42% (6 of 14) requiring increased observation (≥10% weight loss). In contrast, the WT mice held static between days two and three and began to recover by day four, with no mice meeting the requirements for euthanasia and only 13% (2 of 15) requiring increased observation by a margin of ≤0.1g. This Muc4-driven difference in weight loss was not observed in male mice following infection with SARS-CoV (Fig. 2B).

**Fig. 2.**
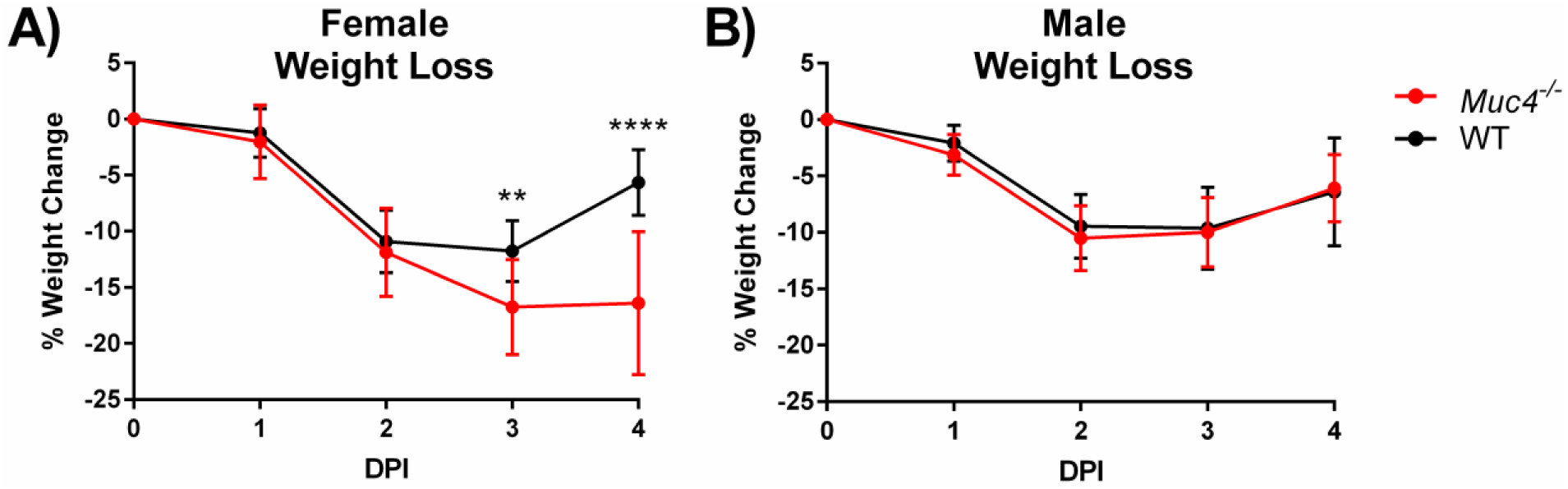
Weight loss following SARS-CoV infection. Weight loss was monitored in (A) female and (B) male *Muc4*^*-*/-^ and WT mice. Black indicates WT mice and red indicates *Muc4*^-/-^ mice. Points represent the mean, and error bars represent standard deviation. Asterisks indicate statistical significance (** = q<0.01, **** = q<0.0001).

### *Muc4* has minimal impact on viral load

Next, the impact of *Muc4*^*-/-*^ on SARS-CoV viral load was measured. Lung lobes from male and female *Muc4*^-/-^ and WT mice were harvested at two DPI, an acute timepoint previously associated with high SARS-CoV titer (Sheahan et al., 2008), and at four DPI, the timepoint at which the QTL on chromosome 16 was mapped (Gralinski et al., 2015). Surprisingly, despite trends for higher mean viral loads in *Muc4*^-/-^ mice on day 2 and day 4, virus titers were not significantly different from WT mice at either timepoint (Fig. 3A). Similarly, despite differences in weight loss, viral titer in the lung also exhibited no sex effect (Table S3). Given the distribution of Muc4 in mucosal surfaces, viral load was further interrogated with IHC to determine whether Muc4 might impact viral tropism in a way not readily determined by the measurement of infectious virus per lobe. Upon examining SARS-CoV antigen staining, however, no significant differences in antigen distribution were noted between *Muc4*^-/-^ and WT mice in either airway or parenchymal staining (Fig. 3B and 3C). Together, the results indicated that the loss of Muc4 had minimal impact on SARS-CoV viral load or distribution.

**Fig. 3.**
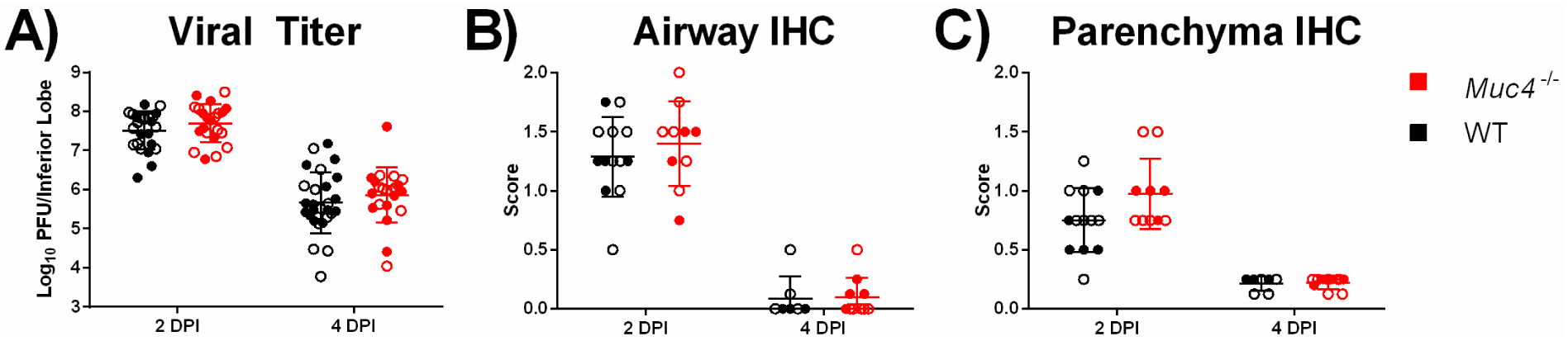
Viral titer and tropism following SARS-CoV infection. (A) viral titer in the inferior lobes of SARS-CoV infected mice at two DPI and four DPI was measured via plaque assay. SARS-CoV tropism in the (B) airway and (C) parenchyma of the left lobe at two and four DPI was measured via blind scoring following IHC staining. Black indicates WT mice and red indicates *Muc4*^-/-^ mice. Points represent individual mice (close = female, open = male), the midline represents the mean, and error bars represent standard deviation. No comparisons of WT versus *Muc4*^-/-^ mice achieved statistical significance after correction for multiple comparisons (q≥0.05)

### *Muc4*^*-/-*^ mice have reduced histopathological damage

Differences in lung damage were assessed using histopathological scoring of H&E stained lungs harvested at either two or four DPI. Similar to weight loss, the histopathology results demonstrated a clear and consistent sex-based phenotype. Female *Muc4*^-/-^ mice had a pattern of lower pathology scores than WT mice (Fig. 4). Airway damage (denudation, debris, and inflammation) was minimally present in female *Muc4*^*-/-*^ at day two or four post-infection (Fig. 4A-C). Similarly, other histopathology and inflammation measures had very low scores in female *Muc4*^-/-^ mice as compared to female WT mice. In contrast, male *Muc4*^*-/-*^ mice induced similar damage to male WT mice (Table S1).

**Fig. 4.**
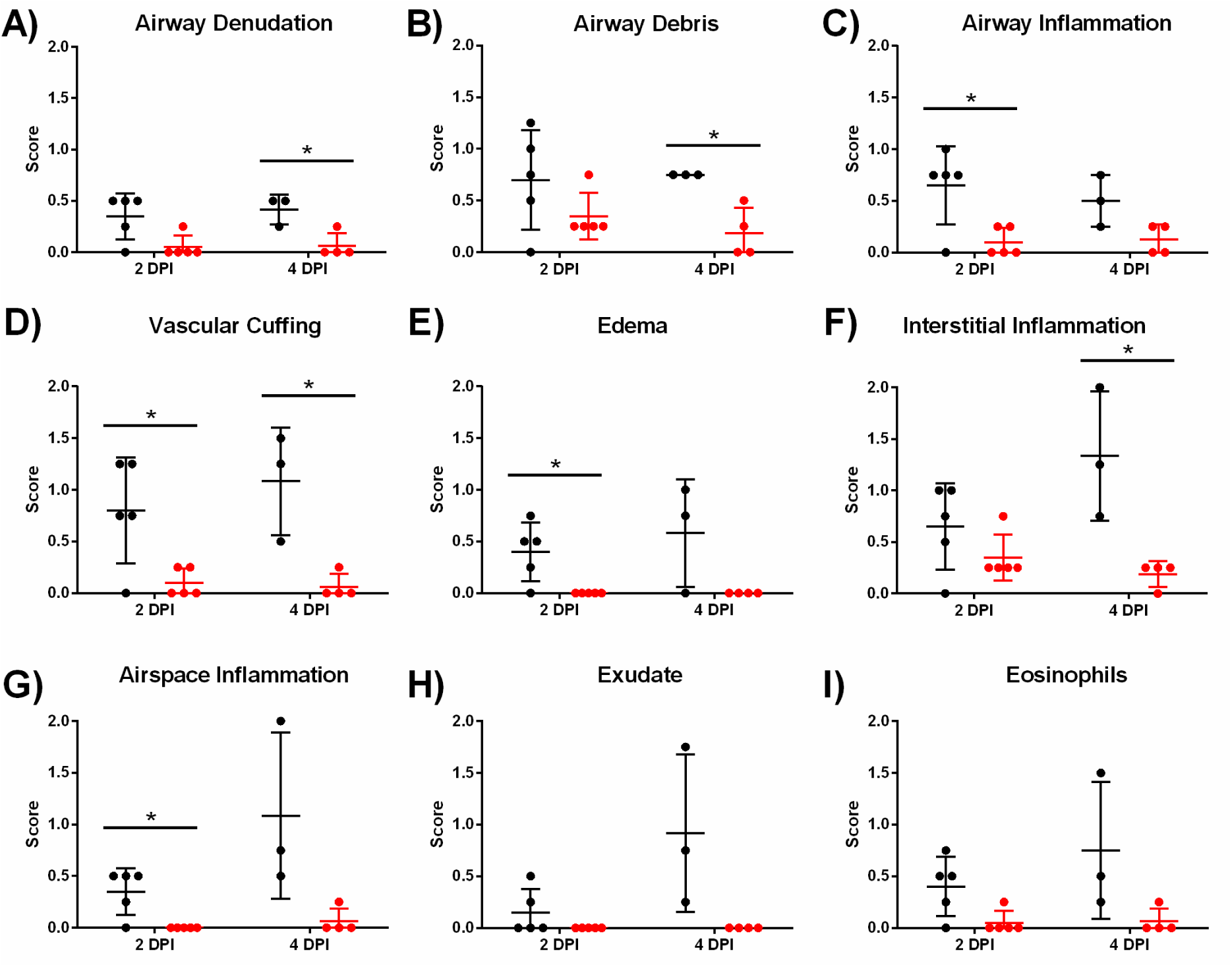
Female WT and *Muc4*^-/-^ mice were infected with SARS-CoV, and their left lung lobes were harvested at either two or four DPI for histopathological analysis following H&E staining. (A) Airway denudation, (B) Airway debris, (C) Airway inflammation, (D) Vascular Cuffing, (E) Edema, (F) Interstitial Inflammation, (G) Airspace Inflammation, (H) Exudate, and (I) Eosinophilia were all scored. Black indicates WT mice and red indicates *Muc4*^-/-^ mice. Points represent individual mice, the midline represents the mean, and error bars represent the standard deviation. Asterisks indicate statistical significance (* = q<0.05).

### Lung function altered in *Muc4*-/- female mice

To further evaluate damage to the lung following infection, whole body plethysmography was utilized to examine changes in pulmonary function (Fig. 5). Using only females, both WT and *Muc4*^-/-^ mice were challenged with a lower dose (10^4^ PFU) to ensure their survival over the full six day time course required to observe the onset, peak, and recovery of disordered lung function following SARS-CoV challenge in mice (Menachery et al., 2015a). Examining airway resistance (penH), the time to peak expiratory flow relative to total expiratory time (rPEF), and mid-tidal expiratory flow (EF50), all three measurements had statistically significant differences between SARS-CoV-infected and mock-infected mice within the *Muc4*^*-/-*^ and WT groups (Supplementary Table S3). However, no Muc4-dependent differences in either penH (Fig. 5A) or rPEF (Fig. 5B) were observed following SARS-CoV infection. In contrast, while not reaching statistical significance, EF50 trended higher in the *Muc4*^*-/-*^ mice as compared to WT mice, peaking at day three post-infection (Fig. 5C). These results show a shift in the early breathing curve in *Muc4*^-/-^ mice with more rapid exhalation and more labored breathing. These findings may have been exacerbated with the original, higher dose, but that experiment was precluded by the limited survival of *Muc4*^-/-^ mice. Overall, the whole body plethysmography data indicate more difficult breathing for female *Muc4*^-/-^ mice relative to their WT counterparts following SARS-CoV infection.

**Fig. 5.**
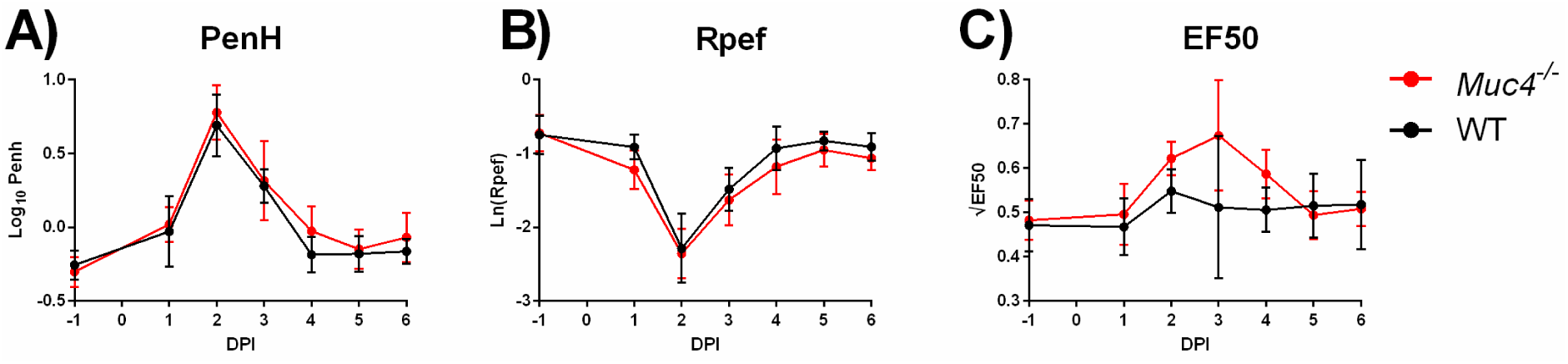
Pulmonary function as measured by normalized (A) PenH, (B) Rpef, and (C) EF50 readings was recorded in SARS-CoV-infected female mice. Black indicates WT mice and red indicates *Muc4*^-/-^ mice. Points represent the mean and error bars represent the standard deviation. Strain-based differences did not achieve statistical significance after correction for multiple comparisons (q≥0.05).

### *Muc4*^*-/-*^ mice have augmented inflammation

Changes in the cytokine and chemokine responses following infection of *Muc4*^*-/-*^ and WT mice were evaluated at two days post infection. While not significant when adjusted for multiple comparisons (Table S3), key inflammatory cytokines including IL-1β, TNF-α, and IL6 had increased expression in *Muc4*^-/-^ mice as compared to WT mice (Fig. 6A-C). Similarly, MIP-1α, MCP-1, and KC also had augmented expression in *Muc4*^-/-^ mice (Fig. 6D-F). Notably, the increased values in *Muc4*^*-/-*^ mice were maintained across both males and females. Together, the results indicate that *Muc4*^*-/-*^ mice have augmented inflammatory cytokine responses relative to WT mice, which may contribute to observed differences in pathogenesis.

**Fig. 6.**
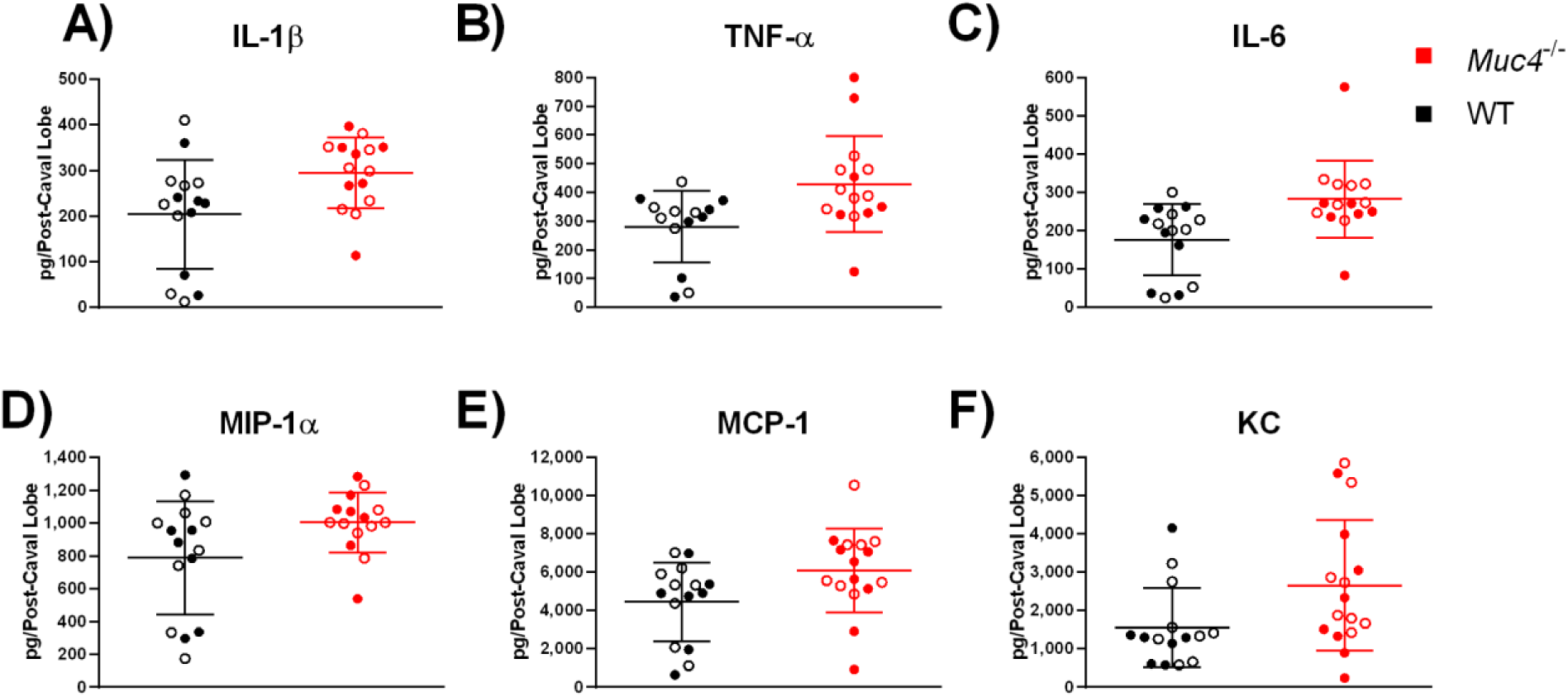
Cytokine levels, including (A) IL-1β, (B) TNF-α, (C) IL-6, (D) MIP-1α, (E) MCP-1, and (F) KC, at 2 DPI in post-caval lobes of SARS-CoV-challenged mice were measured using a BioPlex assay. Black indicates WT mice and red indicates *Muc4*^-/-^ mice. Points represent individual mice (close = female, open = male), the midline represents the mean, and error bars represent standard deviation. Strain-based differences did not achieve statistical significance after correction for multiple comparisons (q≥0.05).

### *Muc4* impacts pathogenesis for an unrelated virus

Because Muc4 modulated SARS-CoV susceptibility independent of viral replication, a cross-platform validation study was undertaken to begin assessing whether the *Muc4* had widespread importance in viral pathogenesis. Chinkungunya virus (CHIKV), an alphavirus, causes inflammatory arthritis and swelling within the joints in human patients. Importantly, *Muc4* has been detected in synovial sarcomas in humans, thus presenting a novel tissue environment and unrelated virus to test the impact of Muc4 (Doyle et al., 2011). Male and female WT and *Muc4*^-/-^ mice were infected with CHIKV and monitored for swelling of the footpad for seven days post infection (Fig. 7). The WT mice have minimal early stage swelling, as was expected with the C57BL/6N model. In contrast, *Muc4*^*-/-*^ mice have augmented disease early during infection, displaying a biphasic swelling pattern. At days 2-4, both male and female mice have increased footpad swelling compared to control mice. This *Muc4*-specific difference was not observed during the second swelling event that took place between days six and seven post-infection. Notably, while sex based differences were observed at days two and four post infection, disease was more robust in male mice, contrasting observations seen with SARS-CoV infection. In either sex, however, the loss of *Muc4* resulted in augmented disease during early time points and indicates a broad role for *Muc4* in viral pathogenesis.

**Fig. 7.**
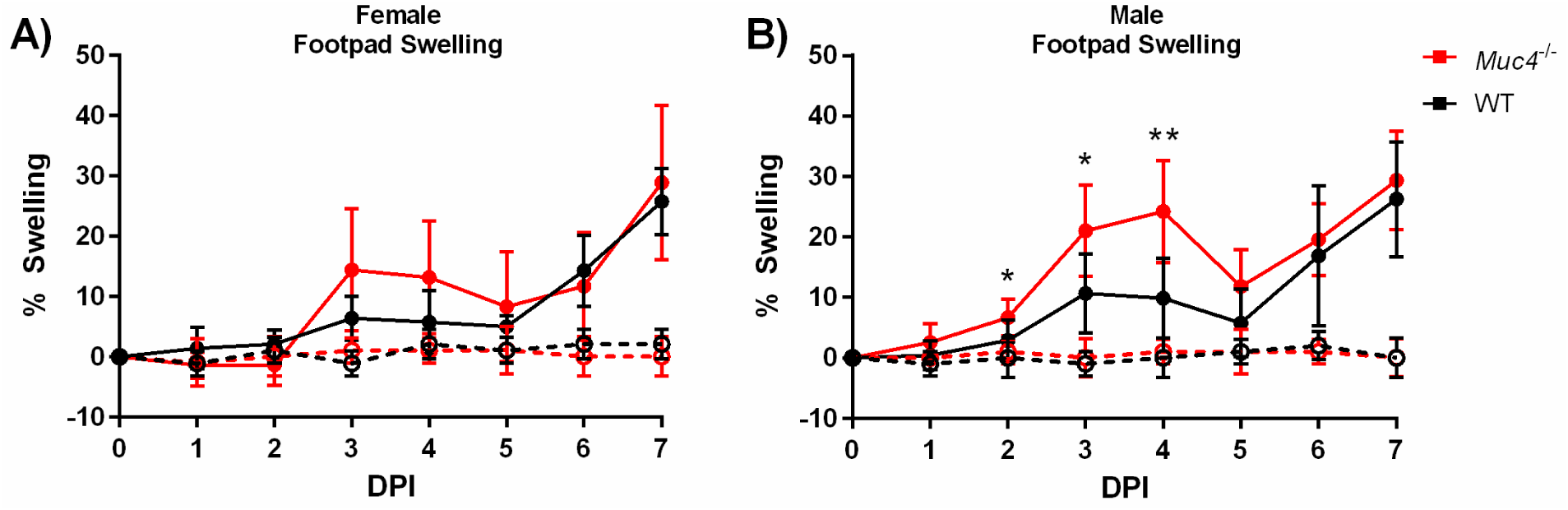
Footpad swelling in (A) female and (B) male mice that have been either mock-infected or infected with CHIKV. Black indicates WT mice and red indicates *Muc4*^-/-^ mice. Dashed lines with open symbols indicate mock infection, and solid lines with solid symbols indicate CHIKV infection. Points represent the mean and error bars represent the standard deviation. Asterisks indicate statistical significance (* = q<0.05, ** = q<0.01).

## Discussion

Screening genetically diverse mouse models provides an opportunity to identify natural variation in novel factors which drive viral disease responses. These studies can also provide therapeutic, prophylactic and molecular insights into emerging pathogens, which are difficult to study during the context of an outbreak. Here, prior phenotypic QTL analysis, bioinformatics, and RNA expression analysis were leveraged to identify *Muc4* as a high priority candidate gene driving differences in SARS-CoV titer. Utilizing a *Muc4* knockout mouse, a role was confirmed for the gene in augmented SARS-CoV pathogenesis. While virus titers trended higher in *Muc4*^-/-^ mice at day 2 and 4, there was no statistically significant change as compared to WT control mice. However, in absolute terms the *Muc4*^-/-^ mice did have a modest 62% higher titer on day two and a 51% higher titer on day four as compared to WT, and it is possible that this difference is biologically significant despite the lack of statistical significance as determined by p-value (Lytsy et al., 2018; Vyas et al., 2015). In addition, the loss of *Muc4* was linked to enhanced CHIKV disease *in vivo*. Together, this study highlights both the utility and the challenges in transitioning from QTL hits to single-molecule studies. Our initial hypothesis for *Muc4* to play a role in controlling virus replication proved incorrect, or, at the very least, substantially more complex than a simple, direct correlate; however, our exploration found a disease-interaction that played a role in pathogenesis across multiple viruses.

Outside of human GWAS studies, where recombination rates are high, it has long proven a challenge to go from a QTL to individual candidate genes (Brown et al., 1997). Even with novel bioinformatic approaches, in the absence of large genomic deletions (Ferris et al., 2013), fortuitously gene-poor regions (Gralinski et al., 2015), or previously identified functions (Gralinski et al., 2017) candidate gene identification continues to be difficult. In this manuscript, we built a framework to integrate knowledge about the preCC genome sequences and a partner microarray dataset to narrow a QTL region down to a likely candidate based on expression correlations. Muc4 exhibited strong correlation data and low expression in the relevant founder strain. Additionally, Muc4 is known to play a role in anti-apoptotic signaling (Chaturvedi et al., 2008; Funes et al., 2006; Moniaux et al., 2007), analysis of published microarray data (Jonckheere N et al., 2012) reveals that cells lacking Muc4 express lower levels of interferon-stimulated genes (Menachery VD, 2014), and the related mucin Muc1 is both anti-apoptotic and anti-inflammatory (Kato K et al., 2014; Li Y et al., 2010; Ueno K et al., 2008). We chose *Muc4* as our priority candidate gene for follow up, with the initial hypothesis that Muc4 suppressed apoptosis and possible the interferon response, and that its absence in a *Muc4*^-/-^ mouse would therefore lead to increased apoptosis and inflammation, thereby inhibiting SARS-CoV replication.

Our validation utilized a *Muc4*^*-/-*^ mouse and confirmed a role for *Muc4* in protection from SARS-CoV- and CHIKV-induced disease and pathogenesis. However, we found essentially no differences in SARS-CoV viral load in the *Muc4*^-/-^ mouse strain compared to WT mice. This may be due to sex (discussed more below), differences between naturally occurring SNPs impacting gene structure and expression and ablative knockouts, or the effects of overall genetic background on given gene variants (Leist et al., 2016). The SARS-CoV titer QTL on chromosome 16 was driven by the PWK/PhJ allele, which was strongly associated with lower viral titer and had low expression of *Muc4*. In contrast, mice with the C56BL/6J allele were associated with high titer and high expression of *Muc4*. Furthermore, PWK/PhJ mice encode functional variants in *Muc4* that are not present in the other seven founder strains. It is therefore possible that C57BL/6 genetic background for the *Muc4*^-/-^ mice, as opposed to a PWK/PhJ background, obfuscated any role for *Muc4* in the regulation of viral titer; this possibility is especially relevant given that the hypothesized role of Muc4 is in the context of complex signaling pathways requiring multiple molecular partners, rather than a direct role such as the physical barrier function of mucins. Alternatively, *Muc4* may simply not impact viral titer following SARS-CoV infection; this leaves the possibility that other genetic targets within the chromosome 16 QTL are responsible for limiting SARS-CoV titers at later time points. Furthermore, the existence of multiple titer-driven QTLs reaffirms that no single gene is expected to serve as the sole driver of SARS-CoV titer. Yet, despite the lack of QTL validation, these studies still identified a role for *Muc4* in viral pathogenesis.

A surprising outcome from these studies was the demonstration of sex-specific and increased disease severity in *Muc4*^-/-^ female mice infected with SARS-CoV. Females lacking Muc4 consistently demonstrated increased susceptibility to SARS-CoV-induced disease, while males did not. These results affirm the National Institute of Health emphasis on conducting animal studies in both male and female subjects (Clayton and Collins, 2014; National Institutes of Health, 2015). Interestingly, SARS-CoV has been reported to cause more severe disease in male mice, with the female mice deriving their resistance at least in part from estrogen signaling (Channappanavar et al., 2017). Furthermore, epidemiological data from both SARS-CoV and SARS-CoV-2 indicate that human females may be more resistant than human males (Chen et al., 2020; Huang et al., 2020; Karlberg et al., 2004; Leong et al., 2006). Estrogen increases *Muc4* transcription in a tissue-specific manner (Lange et al., 2003), offering a potential mechanism as to the greater impact of Muc4’s absence in female mice. Sex effects in the context of *Muc4* have also been reported in a murine model of colitis and colitis-associated colorectal cancer (Das et al., 2016). These results hint that the role of *Muc4* may differ in males and females based upon factors inherent to the host, such as localization, expression levels, or interactions with hormone signaling pathways, and may be less dependent on factors that are unique to a particular disease model.

Notably, a role for Muc4 extended across two different viral pathogens. SARS-CoV infection of female *Muc4*^-/-^ mice, and CHIKV infection of both male and female *Muc4*^-/-^ mice, resulted in more severe disease following infection. For SARS-CoV, loss of Muc4 exacerbated weight loss and increased lethality; surprisingly, less histopathologic damage was observed in the *Muc4*^*-/-*^ lungs following infection despite augmented inflammatory cytokines. The results suggest that the loss of *Muc4* may exacerbate systemic disease and delay the repair/recovery processing in the lung following SARS-CoV infection. A similar reduction in histopathology damage was observed with MERS-CoV infection of immunocompromised rhesus macaques providing evidence for this theory (Prescott et al., 2018). Notably, footpad swelling following CHIKV infection is representative of systemic disease and is exacerbated in the absence of *Muc4.* Although speculative, it is possible that Muc4 functions not to modulate local viral replication, but rather to limit disseminated disease. Clearly, more studies are needed to unravel these complex interactions.

Overall, this research demonstrates that Muc4 is a novel regulator of viral pathogenesis. This study represents the first investigation of Muc4 in the context of any viral infection, and the absence of Muc4 exacerbates disease in two unrelated viruses. The SARS-CoV and CHIKV results indicate that Muc4 has a role in regulating susceptibility and may be widespread across viral pathogens. The study of mucin-driven cell signaling and its role in viral pathogenesis represents a novel avenue of research with the potential to identify new anti-viral therapeutic targets.

## Materials and Methods

### Infection with SARS-CoV or Chikungunya Virus

The generation of C57BL/6NTac *Muc4*^*tm1Unc*^ mice, kindly provided by Dr. Scott Randell of the University of North Carolina at Chapel Hill and hereafter referred to as *Muc4*^*-/-*^ mice, was described by Rowson-Hodel *et al* (Rowson-Hodel et al., 2018). The C57BL/6NTac genetic background of these mice was confirmed in our hands via utilization of the MiniMUGA genotyping array (Neogen Inc, Lincoln, NE), a new genotyping array which includes diagnostic markers for the substrain origin of many inbred mouse strains. Age- and sex-matched wild-type control C57BL/6NTac mice (hereafter referred to as WT mice) were obtained from Taconic (Germantown, NY). Ten- to eleven-week-old mice were anesthetized with a mixture of ketamine (Zoetis, Kalamazoo, MI) and xylazine (Akorn Animal Health, Lake Forest, IL) and inoculated intranasally with either PBS (Gibco, Grand Island, NY), 1×10^4^ PFU, or 1×10^5^ PFU of recombinant mouse adapted SARS-CoV (rMA15) in a 50μl volume (Roberts et al.). Mice receiving the 1×10^5^ PFU dose and their mock-infected counterparts were weighed daily and were euthanized at either two or four days post-infection (DPI). The inferior lobe was harvested and stored intact at −80°C in 1ml PBS with glass beads for downstream titration, and the post-caval lobe was similarly stored for downstream cytokine and chemokine analysis. The left lobe was stored at 4°C in 10% buffered paraformaldehyde (Fisher Scientific, Fair Lawn, NJ) for at least seven days for downstream histopathology and immunohistochemistry. Mice receiving the 1×10^4^ PFU dose and their mock-infected counterparts were weighed daily and their lung function was assessed via whole body plethysmography with in Buxco FinePoint system (Data Sciences International, New Brighton, MN) from one day prior to infection through six DPI. All data from SARS-CoV infections is summarized in Table S1.

Seven-week-old mice were anesthetized with isoflurane (Primal Enterprises, Andrha Pradesh, India) and inoculated subcutaneously in the left rear footpad with either diluent (PBS supplemented with 1% FBS, 1mM CaCl_2_, and 0.5mM MgCl_2_) or 100 PFU of recombinant chikungunya virus (CHIKV) SL15649 in a 10μl volume (Morrison et al., 2011). Footpad size was measured and recorded daily using a caliper to measure the vertical height of the ball of the left rear foot. Mice were humanely euthanized at seven DPI. All data from CHIKV infections is summarized in Table S2.

### Lung Titration

Lung samples were thawed at 37° and lysed for 60 seconds at 6,000 rpm in a MagNA Lyser (Roche, Mannheim, Germany). Debris was pelleted, and lung homogenate was serially diluted 10-fold in PBS. Vero E6 cell monolayers in 6-well plates were infected with 200μl of diluted lung homogenate (10^−1^-10^−6^) for one hour at 37°C with 5% CO_2_. Monolayers were overlaid with a semi-solid overlay containing MEM supplemented with 2% FBS (HyClone, Logan, UT), pencicillin/streptomycin (Gibco, Grand Island, NY), and 0.8% agar (Lonza, Rockland, ME). At two DPI plates were stained with neutral red (Fisher Scientific, Fair Lawn, NJ) for approximately four hours and plaques were visualized with a light box.

### Cytokine Analysis

Cytokine production was measured using the Bio-Plex Pro Mouse Cytokine 23-plex Assay (Bio-Rad, Hercules, CA) on the MAGPIX Multiplex Reader (Bio-Rad, Hercules, CA) according to the manufacturer’s instructions. Lung samples were homogenized and clarified as described for the lung titration assay prior to analysis.

### Histopathology and Immunohistochemistry

Formalin-fixed lobes were embedded in paraffin and 4μm sections were mounted on either Superfrost slides (Fisher Scientific, Waltham, MA) for hematoxylin and eosin staining or on ProbeOn slides (Fisher Scientific, Waltham, MA) for immunohistochemistry (IHC). Embedding, mounting, and hematoxylin and eosin staining was performed by the LCCC Animal Histopathology Core Facility at the University of North Carolina at Chapel Hill.

For IHC, sections were deparaffinized with xylene and rehydrated, then placed in near-boiling antigen retrieval buffer (10mM Tris base, 1mM EDTA, 0.05% Tween-20, pH 9.0) for 20 minutes. Non-specific binding was blocked with 10% normal goat serum (Sigma, St. Louis, MO) and 1% nonfat milk (Lab Scientific, Highlands, NJ) in TBS for two hours at room temperature. Rabbit anti-SARS-CoV nucleocapsid polyclonal antibody (PA1-41098, Thermo Scientific, Rockford, IL) diluted 1:2,000 in blocking buffer was allowed to bind overnight at 4°C. Slides were washed three times in TBS with 0.025% Triton X-100 (TBST) (Sigma, St. Louis, MO), then incubated in 0.3% H_2_O2 in TBS for 15 minutes to prevent endogenous peroxidases from generating a background signal. Slides were incubated with horseradish peroxidase-conjugated goat anti-rabbit IgG secondary antibody (ab97051, Abcam, Cambridge, MA) diluted 1:1,000 in blocking buffer for one hour in a humidity box, then washed three times in TBST. Slides were developed using the Metal Enhanced DAB Substrate Kit (Thermo Scientific, Rockford, IL) and counterstained with Richard-Allan Scientific Modified Mayer’s Hematoxylin (Thermo Scientific, Rockford, IL) and 0.2M lithium carbonate (Sigma, St. Louis, MO) prior to rehydration and mounting. All scoring was conducted in a blinded fashion on a severity scale of 0 (none) to 3 (severe).

### Statistical Analysis

RNA expression levels were analyzed by one-way ANOVA with Dunnett’s multiple comparisons test comparing PWK/PhJ to the other seven CC founder lines using Prism 7 for Windows version 7.01 (GraphPad Software, La Jolla, CA). Buxco data was transformed as previously described (Menachery et al., 2015a) Daily results for all titer-related data types (log_10_-transformed viral load and immunohistochemistry) and cytokine levels were analyzed by two-way ANOVA using RStudio version 1.1.383 with R version 3.2.0 (R Core Team, 2017; RStudio Team, 2015) to determine the impact of mouse strain and sex. Transformed daily Buxco results were similarly analyzed by two-way ANOVA to determine the impact of infection and mouse strain. Daily results for all pathology-related data types (histopathology, weight change, and footpad swelling) was analyzed by three-way ANOVA to determine the impact of mouse strain, sex, and infection status. When three-way ANOVA results indicated significant differences on the basis of sex (weight loss, histopathology, and footpad swelling), the strain-based differences between infected animals of a single sex were analyzed using multiple t-tests with the Benjamini and Hochberg correction for false discoveries in Prism. All graphing was done in Prism. A significance threshold of >0.05 after correction for multiple comparisons was considered significant throughout all studies. The results of all statistical analyses are summarized in Table S3.

### Data Availability

All phenotypic data associated with the infection of *Muc4*^-/-^ mice and their WT counterparts are summarized in Table S1 and S2. *Muc4*^-/-^ mice are available upon request.

## Author Contributions

JAP was involved with the design of all experiments, performed all SARS-CoV mouse experiments, and prepared the manuscript. KSP and MTH designed and performed the CHIKV mouse experiments. LEG provided feedback regarding experimental design and performed SARS-CoV mouse experiments and BioPlex analysis. AB performed immunohistochemistry staining. VDM provided feedback regarding experimental design, assisted with SARS-CoV mouse experiments, and assisted substantially with the editing of the manuscript. MTF provided guidance regarding statistical analysis. MTF, DB, and SKM performed array analysis and assisted with the selection of *Muc4* from the list of potential target genes. RSB scored gross pathology and immunohistochemistry and contributed to all aspects of study design, experimental design, and manuscript design and editing.

## Acknowledgements

The authors thank Dr. Scott H. Randell of the Department of Cell Biology and Physiology at the University of North Carolina at Chapel Hill for contributing the *Muc4*^*-/-*^ mice, without which these studies would not have been possible.

